# A simple method for automated equilibration detection in molecular simulations

**DOI:** 10.1101/021659

**Authors:** John D. Chodera

**Affiliations:** Computational Biology Program, Sloan Kettering Institute, Memorial Sloan Kettering Cancer Center, New York, NY 10065

**Keywords:** molecular dynamics (MD), Metropolis-Hastings, Monte Carlo (MC), Markov chain Monte Carlo (MCMC), equilibration, burn-in, timeseries analysis, statistical inefficiency, integrated autocorrelation time

## Abstract

Molecular simulations intended to compute equilibrium properties are often initiated from configurations that are highly atypical of equilibrium samples, a practice which can generate a distinct initial transient in mechanical observables computed from the simulation trajectory. Traditional practice in simulation data analysis recommends this initial portion be discarded to *equilibration,* but no simple, general, and automated procedure for this process exists. Here, we suggest a conceptually simple automated procedure that does not make strict assumptions about the distribution of the observable of interest, in which the equilibration time is chosen to maximize the number of effectively uncorrelated samples in the production timespan used to compute equilibrium averages. We present a simple Python reference implementation of this procedure, and demonstrate its utility on typical molecular simulation data.

## INTRODUCTION

Molecular simulations use Markov chain Monte Carlo (MCMC) techniques [1] to sample configurations *x* from an equilibrium distribution *π*(*x*), either exactly (using Monte Carlo methods such as Metropolis-Hastings) or approximately (using molecular dynamics integrators without Metropolization) [2].

Due to the sensitivity of the equilibrium probability density *π*(*x*) to small perturbations in configuration *x* and the difficulty of producing sufficiently good guesses of typical equilibrium configurations *x ~ π*(*x*), these molecular simulations are often started from highly atypical initial conditions. For example, simulations of biopolymers might be initiated from a fully extended conformation unrepresentative of behavior in solution, or a geometry derived from a fit to diffraction data collected from a cryocooled crystal; solvated systems may be prepared by periodically replicating a small solvent box equilibrated under different conditions, yielding atypical densities and solvent structure; liquid mixtures or lipid bilayers may be constructed by using methods that fulfill spatial constraints (e.g. PackMol [3]) but create locally aytpical geometries, requiring long simulation times to relax to typical configurations.

As a result, traditional practice in molecular simulation has recommended some initial portion of the trajectory be discarded to *equilibration* (also called *burn-in*^1^ in the MCMC literature [4]). While the process of discarding initial samples is strictly unnecessary for the time-average of quantities of interest to eventually converge to the desired expectations [5], this nevertheless often allows the practitioner to avoid what may be impractically long run times to eliminate the bias in computed properties in finite-length simulations induced by atypical initial starting conditions. It is worth noting that a similar procedure is not a practice universally recommended by statisticians when sampling from posterior distributions in statistical inference [4]; the differences in complexity of probability densities typically encountered in statistics and molecular simulation may explain the difference in historical practice.

As a motivating example, consider the computation of the average density of liquid argon under a given set of reduced temperature and pressure conditions shown in Figure 1. To initiate the simulation, an initial dense liquid geometry at reduced density *ρ*^*^ = *ρσ*^3^ = 0.960 was prepared and subjected to local energy minimization. The upper panel of Figure 1 depicts the average relaxation behavior of simulations initiated from the same configuration with different random initial velocities and integrator random number seeds (see *Simulation Details*). The average of 500 realizations of this process shows a characteristic relaxation away from the initial density toward the equilibrium density (Figure 1, upper panel, black line). As a result, the expectation of the running average of the density significantly deviates from the true expectation (Figure 1, lower panel, dashed line). This effect leads to significantly biased estimates of the expectation unless simulations are sufficiently long to eliminate starting point dependent bias, which takes a surprisingly long ~30 ns in this example. Note that this bias is present even in the average of many realizations because the *same* atypical starting condition is used for every realization of this simulation process.

**FIG. 1.**
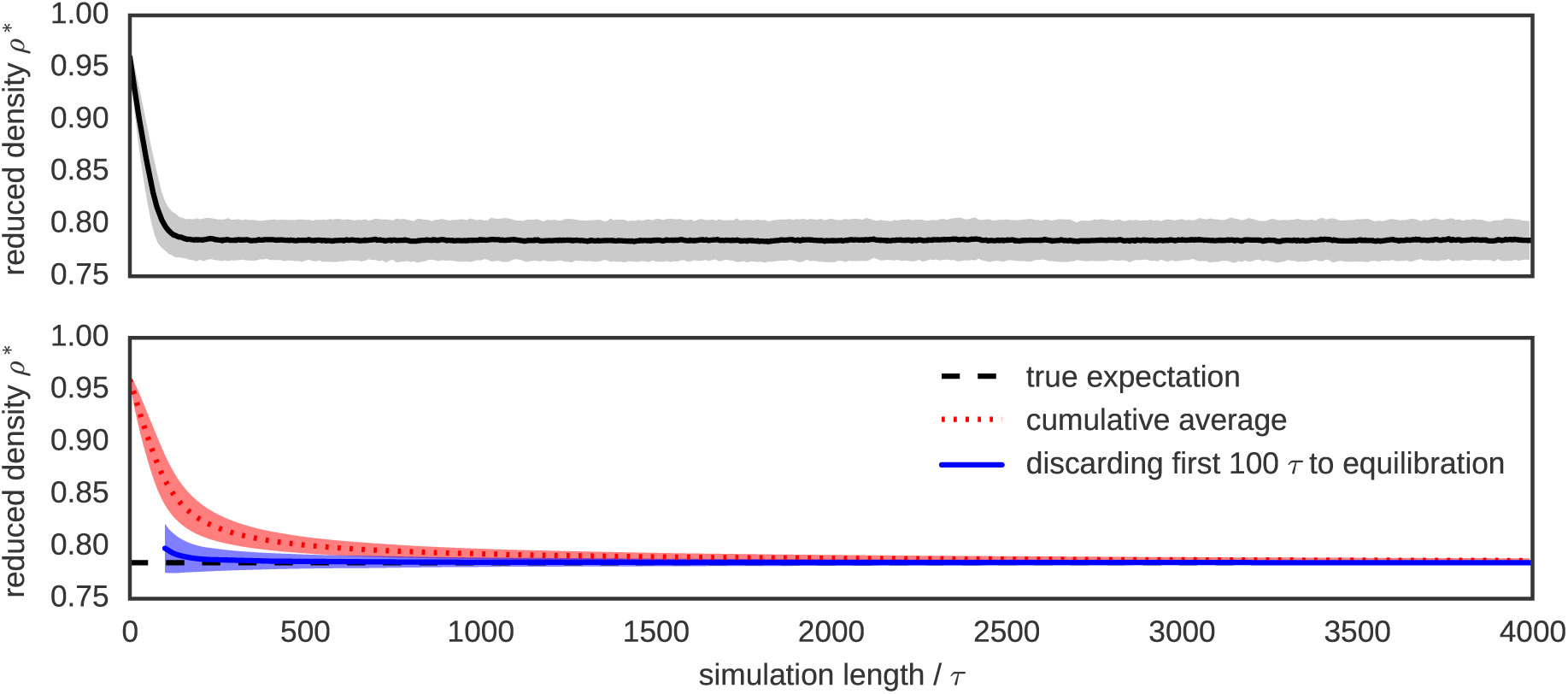
Illustration of the motivation for discarding data to equilibration. To illustrate the bias in expectations induced by relaxation away from initial conditions, 500 replicates of a simulation of liquid argon were initiated from the same energy-minimized initial configuration constructed with initial reduced density *ρ*^*^ = *ρσ*^3^ = 0.960 but different random number seeds for stochastic integration. Top: The average of the reduced density (black line) over the replicates relaxes to the region of typical equilibrium densities over the first ~ 100*τ* of simulation time, where *τ* is a natural time unit (see *Simulation Details*). Bottom: If the average density is estimated by a cumulative average from the beginning of the simulation (red dotted line), the estimate will be heavily biased by the atypical starting density even beyond 1000*τ*. Discarding even a small amount of initial data—in this case 500 initial samples—results in a cumulative average estimate that converges to the true average (black dashed line) much more rapidly. Shaded regions denote 95% confidence intervals.

To develop an automatic approach to eliminating this bias, we take motivation from the concept of *reverse cumulative averaging* from Yang et al. [6], in which the trajectory statistics over the production region of the trajectory are examined for different choices of the end of the discarded equilibration region to determine the optimal production region to use for computing expectations and other statistical properties. We begin by first formalizing our objectives mathematically.

Consider *T* successively sampled configurations *x_t_* from a molecular simulation, with *t* = 1,…, *T*, initiated from *x*_0_. We presume we are interested in computing the expectation

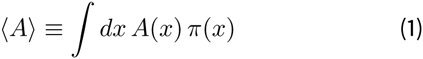

of a mechanical property of interest A(*x*). For convenience, we will refer to the timeseries *a_t_* ≡ *A*(*x_t_*), with *t ϵ* [1, *T*]. The estimator Â ≈ 〈A〉 constructed from the entire dataset is given by

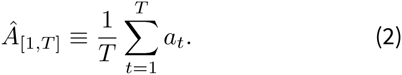

While lim*_T→∞_ Â*_[1,_*^T^*_]_ = 〈A〉 for an infinitely long simulation^2^, the bias in *Â*_[1,_*^T^*_]_ may be significant in a simulation of finite length *T*.

By discarding samples *t* < *t*_0_ to equilibration, we hope to exclude the initial transient from our sample average, and provide a less biased estimate of 〈A〉,

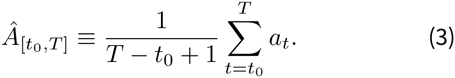

We can quantify the overall error in an estimator 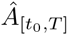 in a sample average that starts at *x*_0_ and excludes samples where *t* < *t*_0_ by the expected error 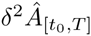

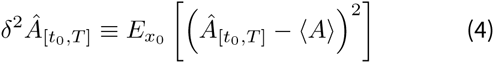

where 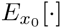denotes the expectation over independent realizations of the specific simulation process initiated from configuration *x*_0_, but with different velocities and random number seeds.

We can rewrite the expected error *δ*^2^*Â* by separating it into two components:

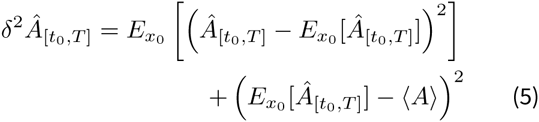

The first term denotes the variance in the estimator *Â*,

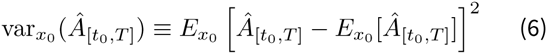

while the second term denotes the contribution from the squared bias,

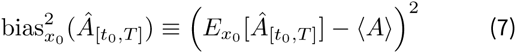

With increasing equilibration time *t*_0_, bias is reduced, but the variance—the contribution to error due to random variation from having a finite number of uncorrelated samples—will increase because less data is included in the estimate. This can be seen in the bottom panel of Figure 2, where the shaded region (95% confidence interval of the mean) increases in width with increasing equilibration time *t*_0_.

**FIG. 2.**
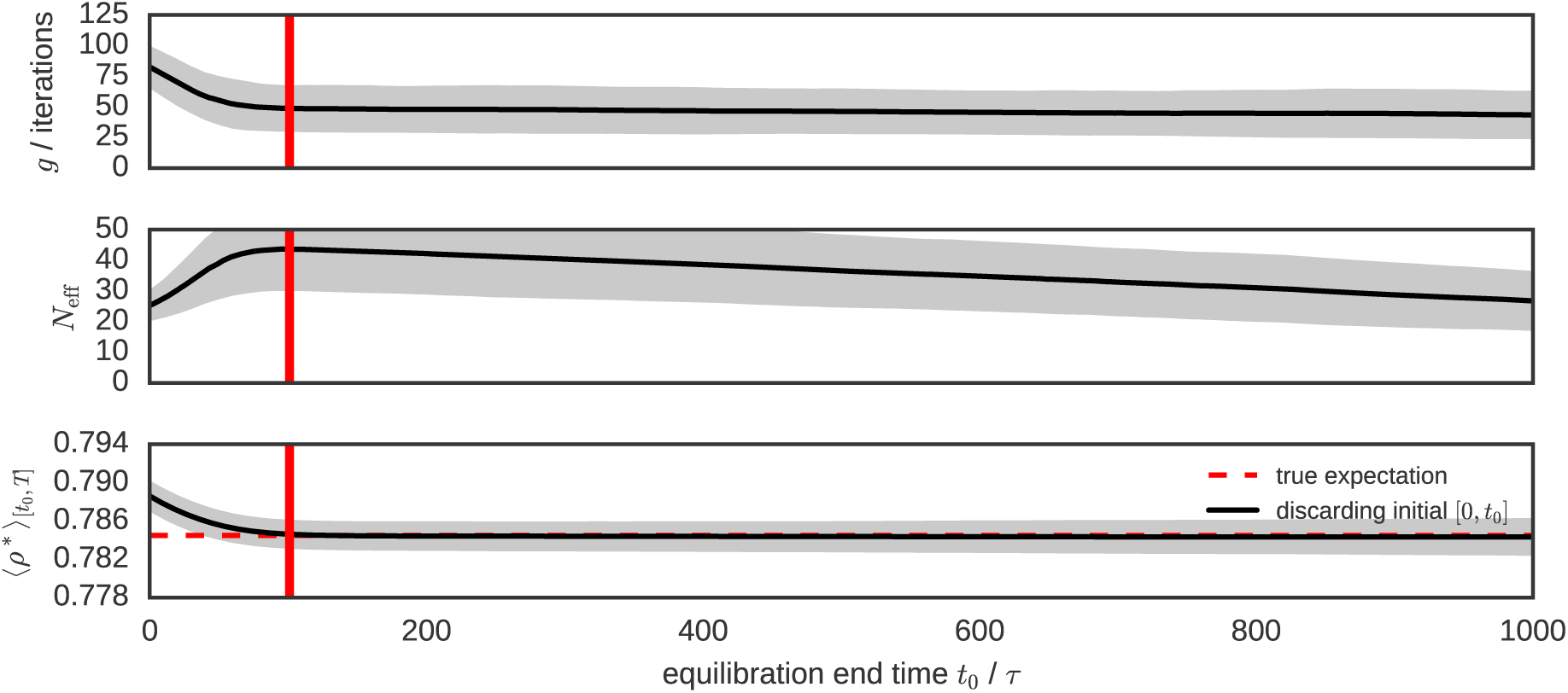
Statistical inefficiency, number of uncorrelated samples, and bias for different equilibration times. Trajectories of length *T* = 2000*τ* for the argon system described in were analyzed as a function of equilibration time choice *t*_0_. Averages over all 500 replicate simulations (all starting from the same initial conditions) are shown as dark lines, with shaded lines showing standard deviation of estimates among replicates. **Top**: The statistical inefficiency *g* as a function of equilibration time choice *t*_0_ is initially very large, but diminishes rapidly after the system has relaxed to equilibrium. **Middle**: The number of effectively uncorrelated samples *N*_eff_ = (*T* − *t*_0_ + 1)/*g* shows a maximum at *t*_0_ ~ 100*τ* (red vertical lines), suggesting the system has equilibrated by this time. **Bottom**: The cumulative average density 〈*p\**〉 computed over the span [to, *T*] shows that the bias (deviation from the true estimate, shown as red dashed lines) is minimized for choices of *t*_0_ ≥ 100*τ*. The standard deviation among replicates (shaded region) grows with *t*_0_ because fewer data are included in the estimate. The choice of optimal *t*_0_ that maximizes *N*_eff_ (red vertical line) strikes a good balance between bias and variance. The true estimate (red dashed lines) is computed from averaging over the range [5 000, 10 000]*τ* over all 500 replicates.

To examine the tradeoff between bias and variance explicitly, Figure 3 plots the bias and variance (here, shown as standard error) contributions against each other as a function of *t*_0_ (denoted by color) as computed from statistics over all 500 replicates. At *t*_0_ = 0, the bias is large but variance is minimized. With increasing *t*_0_, bias is eventually eliminated but then variance rapidly grows as fewer uncorrelated samples are included in the estimate. There is a clear optimal choice at *t*_0_ ~ 100*τ* that minimizes variance while also effectively eliminating bias (where *τ* is a natural time unit—see *Simulation Details*).

**FIG. 3.**
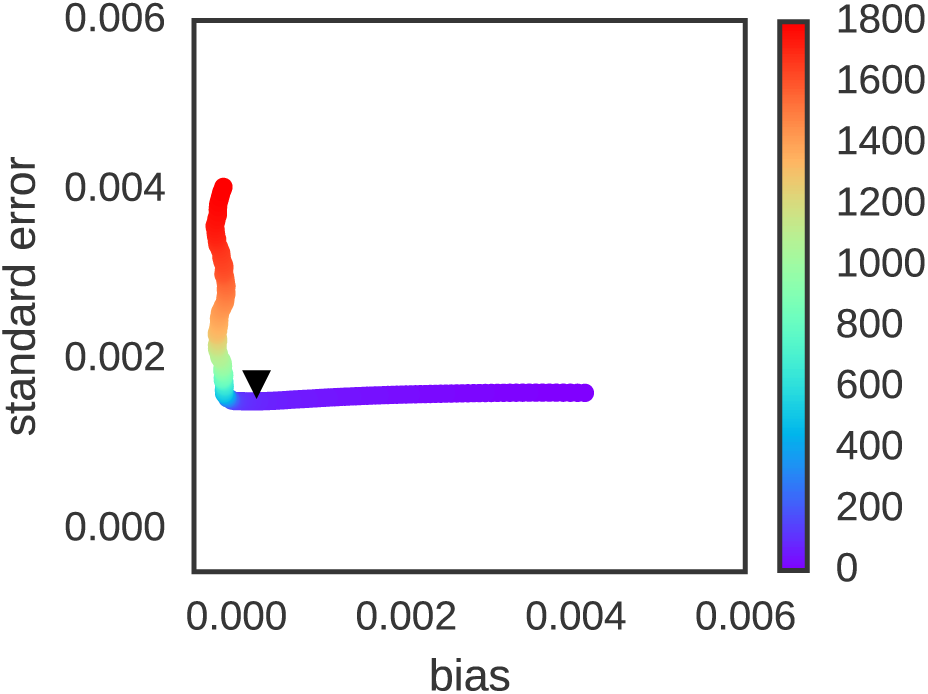
Bias-variance tradeoff for fixed equilibration time versus automatic equilibration time selection. Trajectories of length *T* = 2000*τ* for the argon system described in were analyzed as a function of equilibration time choice *t*_0_, with colors denoting the value of *t*_0_ (in units of *τ*) corresponding to each plotted point. Using 500 replicate simulations, the average bias (average deviation from true expectation) and standard deviation (random variation from replicate to replicate) were computed as a function of a prespecified fixed equilibration time *t*_0_, with colors running from violet (0 *τ*) to red (1800 *τ*). As is readily discerned, the bias for small *t*_0_ is initially large, but minimized for larger *t*_0_. By contrast, the standard error (a measure of variance, estimated here by standard deviation among replicates) grows as *t*_0_ grows above a certain critical time (here, ~ 100*τ*). If the *t*_0_ that maximizes *N*_eff_ is instead chosen *individually* for each trajectory based on that trajectory’s estimates of statistical inefficiency 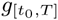, the resulting bias-variance tradeoff (black triangle) does an excellent job minimizing bias and variance simultaneously, comparable to what is possible for a choice of equilibration time *t*_0_ based on knowledge of the true bias and variance among many replicate estimates.

## SELECTING THE EQUILIBRATION TIME

Is there a simple approach to choosing an optimal equilibration time *t*_0_ that provides a significantly improved estimate 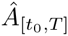 even when we do not have access to multiple realizations? At worst, we hope that such a procedure would at least give some improvement over the naive estimate, such that 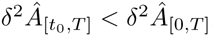; at best, we hope that we can achieve a reasonable bias-variance tradeoff close to the optimal point identified in Figure 3 that minimizes bias without greatly increasing variance. We remark that, for cases in which the simulation is not long enough to reach equilibrium, no choice of *t*_0_ will eliminate bias completely; the best we can hope for is to minimize this bias.

While automated methods for selecting the equilibration time *t*_0_ have been proposed, these approaches have shortcomings that have greatly limited their use. The reverse cumulative averaging (RCA) method proposed by Yang et al. [6], for example, uses a statistical test for normality to determine the point before which which the observable timeseries deviates from normality when examining the timeseries in reverse. While this concept may be reasonable for experimental data, where measurements often represent the sum of many random variables such that the central limit theorem’s guarantee of asymptotic normality ensures the distribution of the observable will be approximately normal, there is no such guarantee that instantaneous measurements of a simulation property of interest will be normally distributed. In fact, many properties will be decidedly *non-normal.* For a biomolecule such as a protein, for example, the radius of gyration, end-to-end distance, and torsion angles sampled during a simulation will all be highly nonnormal. Instead, we require a method that makes no assumptions about the nature of the distribution of the property understudy.

## AUTOCORRELATION ANALYSIS

The set of successively sampled configurations {*x_t_*} and their corresponding observables {*a_t_*} compose a correlated timeseries of observations. To estimate the statistical error or uncertainty in a stationary timeseries free of bias, we must be able to quantify the *effective number of uncorrelated samples* present in the dataset. This is usually accomplished through computation of the *statistical inefficiency g*, which quantifies the number of correlated time-series samples needed to produce a single effectively uncorrelated sample of the observable of interest. While these concepts are well-established for the analysis of both Monte Carlo and molecular dynamics simulations [7–10], we review them here for the sake of clarity.

For a given equilibration time choice *t*_0_, the statistical uncertainty in our estimator 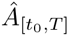 can be written as,

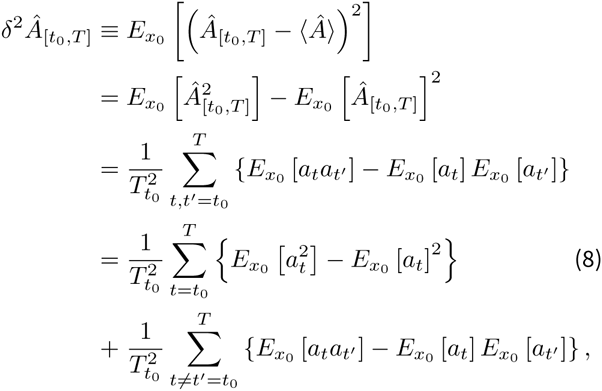

where 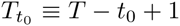 the number of correlated samples in the timeseries 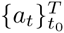. In the last step, we have split the double-sum into two separate sums—a term capturing the variance in the observations *a_t_*, and a remaining term capturing the correlation between observations.

If *t*_0_ is sufficiently large for the initial bias to be eliminated, the remaining timeseries 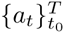 will obey the properties of both *stationarity* and *time-reversibility,* allowing us to write,

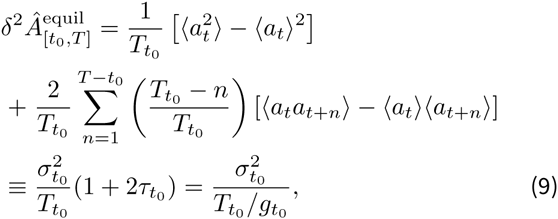

where the variance *σ*^2^, statistical inefficiency *g*, and integrated autocorrelation time *τ* (in units of the sampling interval) are given by

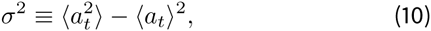

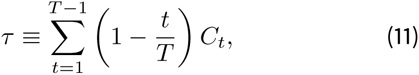

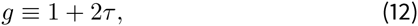

with the discrete-time normalized fluctuation autocorrelation function *C_t_* defined as

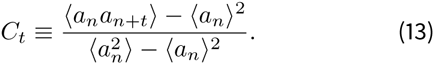

In practice, it is difficult to estimate *C_t_* for *t ~ T*, due to growth in the statistical error, so common estimators of *g* make use of several additional properties of *C_t_* to provide useful estimates (see *Practical Computation of Statistical Inefficiencies*).

The *t*_0_ subscript for the variance *σ*^2^, the integrated auto-correlation time *τ*, and the statistical inefficiency *t*_0_ mean that these quantities are only estimated over the production portion of the timeseries, 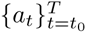 Since we assumed that the bias was eliminated by judicious choice of the equilibration time *t*_0_, this estimate of the statistical error will be poor for choices of *t*_0_ that are too small.

## THE ESSENTIAL IDEA

Suppose we choose some arbitrary time *t*_0_ and discard all samples *t ϵ* [0, *t*_0_) to equilibration, keeping [*t*_0_, *T*] as the dataset to analyze. How much data remains? We can determine this by computing the statistical inefficiency 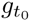 for the interval [*t*_0_, *T*], and computing the effective number of uncorrelated samples 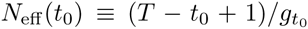 If we start at *t*_0_ ≡ *T* and move *t*_0_ to earlier and earlier points in time, we expect that the effective number of uncorrelated samples *N*_eff_ (*t*_0_) will continue to grow until we start to include the highly atypical initial data. At that point, the integrated autocorrelation time *τ* (and hence the statistical inefficiency *g*) will greatly increase (a phenomenon observed earlier, e.g. Figure 2 of [6]). As a result, the effective number of samples *N*_eff_ will start to plummet.

Figure 2 demonstrates this behavior for the liquid argon system described above, using averages of the statistical inefficiency 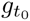 and *N*_eff_ (*t*_0_) computed over 500 independent replicate trajectories. At short *t*_0_, the average statistical inefficiency *g* (Figure 2, top panel) is large due to the contribution from slow relaxation from atypical initial conditions, while at long *t*_0_ the statistical inefficiency estimate is much shorter and nearly constant of a large span of time origins. As a result, the average effective number of uncorrelated samples *N*_eff_ (Figure 2, middle panel) has a peak at *t*_0_ ~ 100*τ* (Figure 2, vertical red lines). The effect on bias in the estimated average reduced density 〈p^*^〉 (Figure 2, bottom panel) is striking—the bias is essentially eliminated for the choice of equilibration time *t*_0_ that maximizes the number of uncorrelated samples *N*_eff_.

This suggests an alluringly simple algorithm for identifying the optimal equilibration time—pick the *t*_0_ which maximizes the number of uncorrelated samples *N*_eff_ in the timeseries 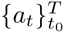 for the quantity of interest *A*(*x*):

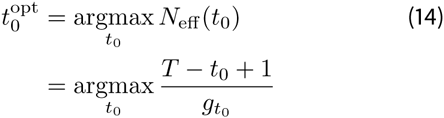

*Bias-variance tradeoff.* How will the simple strategy of selecting the equilibration time *t*_0_ using Eq 14 work for cases where we do not know the statistical inefficiency *g* as a function of the equilibration time *t*_0_ precisely? When all that is available is a single simulation, our best estimate of 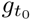 is derived from that simulation alone over the span [*t*_0_, *T*] — will this affect the quality of our estimate of equilibration time? Empirically, this does not appear to be the case—the black triangle in Figure 3 shows the bias and variance contributions to the error for estimates computed over the 500 replicates where *t*_0_ is individually determined from each simulation using this simple scheme based on selecting *t*_0_ to maximize *N*_eff_ for each individual realization. Despite not having knowledge about multiple realizations, this strategy effectively achieves a near-optimal balance between minimizing bias without increasing variance.

*Overall RMS error.* How well does this strategy perform in terms of decreasing the *overall* error 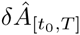 compared to 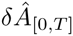? Figure 4 compares the expected standard error (denoted *δÂ*) as a function of a fixed initial equilibration time *t*_0_ (black line with shaded region denoting 95% confidence interval) with the strategy of selecting *t*_0_ to maximize *N*_eff_ for each realization (red line with shaded region denoting 95% confidence interval). While the minimum error for the fixed *t*_0_ strategy (0.00152±0.00005) is achieved at ~ 100*τ*—a fact, that could only be determined from knowledge of multiple realizations—the simple strategy of selecting *t*_0_ using Eq. 14 achieves a minimum error of 0.00173±0.00005, only 11% worse (compared to errors of 0.00441±0.00007, or 290% worse, should no data have been discarded).

**FIG. 4.**
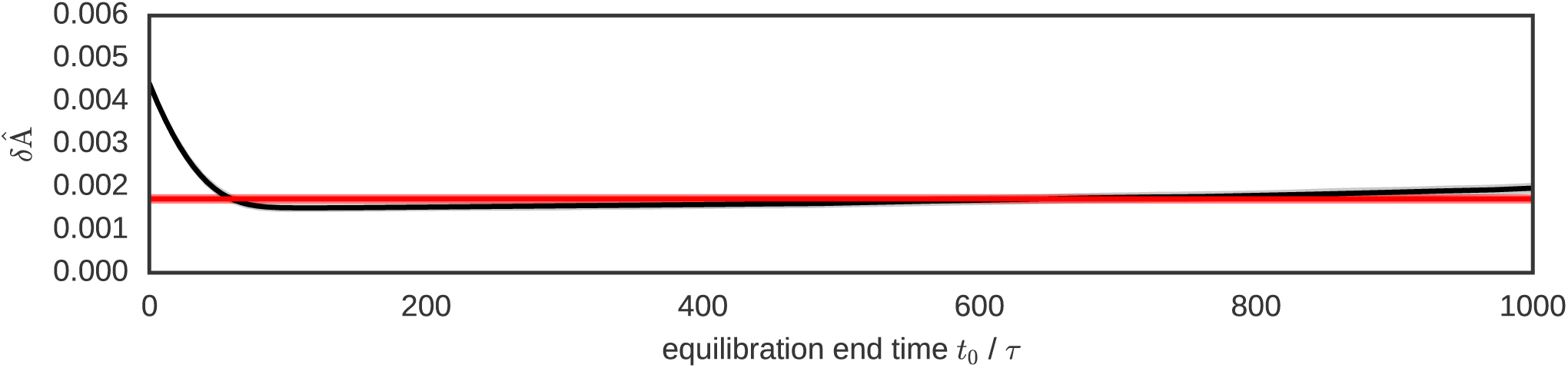
RMS error for fixed equilibration time versus automatic equilibration time selection. Trajectories of length *T* = 2000*τ* for the argon system described in were analyzed as a function of fixed equilibration time choice *t*_0_. Using 500 replicate simulations, the root-mean-squared (RMS) error (Eq. 4) was computed (black line) along with 95% confidence interval (gray shading). The RMS error is minimized for fixed equilibration time choices in the range 100–200*τ*. If the *t*_0_ that maximizes *N*_eff_ is instead chosen *individually* for each trajectory based on that trajectory’s estimated statistical inefficiency 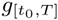 using Eq. 14, the resulting RMS error (red line, 95% confidence interval shown as red shading) is quite close to the minimum RMS error achieved from any particular *fixed* choice of equilibration time *t*_0_, suggesting that this simple automated approach to selecting *t*_0_ achieves close to optimal performance.

## DISCUSSION

The scheme described here—in which the equilibration time *t*_0_ is computed using Eq. 14 as the choice that maximizes the number of uncorrelated samples in the production region [*t*_0_, *T*]—is both conceptually and computation, ally straightforward. It provides an approach to determining the optimal amount of initial data to discard to equilibration in order to minimize variance while also minimizing initial bias, and does this without employing statistical tests that require generally unsatisfiable assumptions of normality of the observable of interest. All that is needed is to save the timeseries 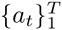 of the observable *A*(*x*) of interest—there is no need to store full configurations *x_t_*—and postprocess this dataset with a simple analysis procedure, for which we have provided a convenient Python reference implementation (see *Simulation Details*). As we have seen, this scheme empirically appears to select a practical compromise between bias and variance even when the statistical inefficiency *g* is estimated directly from the trajectory using Eq. 12.

To show that this approach is indeed general, we repeated the analysis illustrated above in Figs. 1–4 for a different choice of observable *A*(*x*) for the same liquid argon system—in this case, the reduced potential energy^3^ *u^*^*(*x*) *≡βU*(*x*). The results of this analysis are collected in Fig. 5. As, can readily be seen, this reduced potential behaves in essentially the same way the reduced density does, and the simple scheme for automated determination of equilibration time *t*_0_ from Eq. 14 does just as well.

**FIG. 5.**
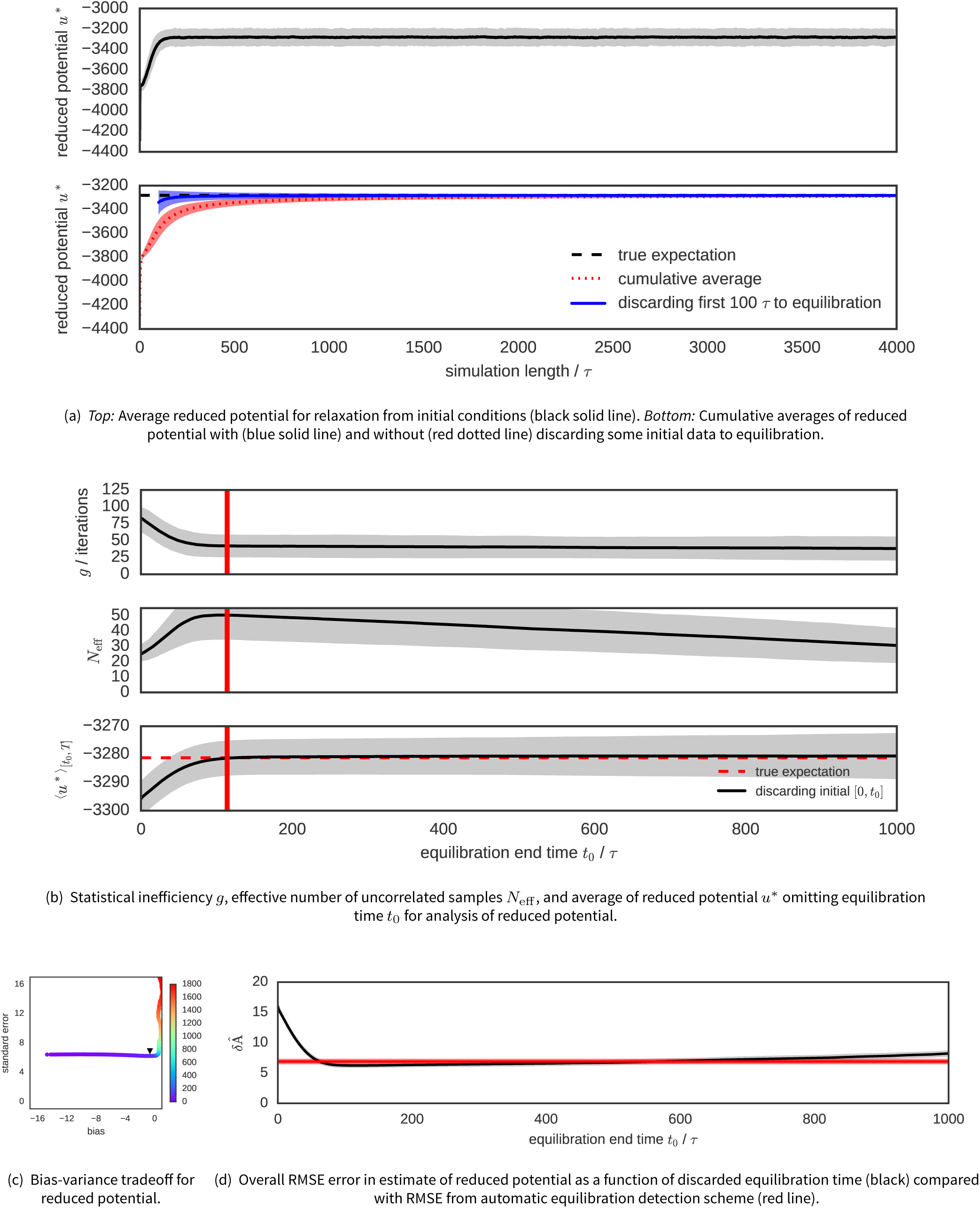
Corresponding analysis for reduced potential energy of liquid argon system. The analyses of Figs. 1–4 were repeated for the reduced potential energy *u*\*(*x*) = *βU* (x) of the liquid argon system. As with the analysis of reduced density, the simple automated determination of equilibration time *t*_0_ from Eq. 14 works equivalently well for the reduced potential. Shaded regions denote 95% confidence interval.

Aword of caution is necessary. One can certainly envision pathological scenarios where this algorithm for selecting an optimal equilibration time will break down. In cases where the simulation is not long enough to reach equilibrium—let alone collect many uncorrelated samples from it—no choice of equilibration time will bestow upon the experimenter the ability to produce an unbiased estimate of the true expectation. Similarly, in cases where insufficient data is available for the statistical inefficiency to be estimated well, this algorithm is expected to perform poorly. However, in these cases, the data itself should be suspect if the trajectory is not at least an order of magnitude longer than the minimum estimated autocorrelation time.

## SIMULATION DETAILS

All molecular dynamics simulations described here were performed with OpenMM 6.3 [12] (available at openmm.org) using the Python API. All scripts used to retrieve the software versions used here, run the simulations, analyze data, and generate plots—along with the simulation data itself and scripts for generating figures—are available on GitHub^4^.

To model liquid argon, the `LennardJonesFluid` model system in the openmmtools package^5^ was used with parameters appropriate for liquid argon (*σ* = 3.4 Å, ϵ = 0.238 kcal/mol). All results are reported in reduced (dimensionless) units. A cubic switching function was employed, with the potential gently switched to zero over *r* ∈ [*σ*, 3*σ*], and a long-range isotropic dispersion correction accounting for this switching behavior used to include neglected contributions. Simulations were performed using a periodic box of *N* = 500 atoms at reduced temperature *T*\* = *k^B^T*/ϵ = 0.850 and reduced pressure *p* ≡ pσ*^3^*/ϵ* = 1.266 using a Langevin integrator [13] with timestep Δ*t* = 0.01*τ* and collision rate *v = τ^−1^*, with characteristic oscillation timescale 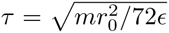 and *r*_0_ = 2^1/6^*σ* [14]. All times are reported in multiples of the characteristic timescale τ. A molecular scaling Metropolis Monte Carlo barostat with Gaussian simulation volume change proposal moves attempted every *τ* (100 timesteps), using an adaptive algorithm that adjusts the proposal width during the initial part of the simulation [12]. Densities were recorded every *τ* (100 timesteps). The true expectation 〈*ρ*\*〉 was estimated from the sample average over all 500 realizations over [5000,10000] *τ*.

The automated equilibration detection scheme is also available in the `timeseries` module of the `pymbar` package as `detectEquilibration()`, and can be accessed using the following code:

~~~
from pymbar.timeseries import detectEquilibration
*# determine equilibrated region*
[*t*0, g, Neff_max] = detectEquilibration(A_t)
*# discard initial samples to equilibration*
A_t = A_t [t0:]
~~~

## PRACTICAL COMPUTATION OF STATISTICAL INEFFICIENCIES

The robust computation of the statistical inefficiency *g* (defined by Eq. 12) for a finite timeseries *a_t_, t* = 0*,…, T* deserves some comment. There are, in fact, a variety of schemes for estimating *g* described in the literature, and their behaviors for finite datasets may differ, leading to different estimates of the equilibration time *t*_0_ using the algorithm of Eq. 14.

The main issue is that a straightforward approach to estimating the statistical inefficiency using Eqs. 11–13 in which the expectations are simply replaced with sample estimates causes the statistical error in the estimated correlation function *C_t_* to grow with *t* in a manner that allows this error to quickly overwhelm the sum of Eq. 11. As a result, a number of alternative schemes—generally based on controlling the error in the estimated *C_t_* or truncating the sum of Eq. 11 when the error grows too large—have been proposed.

For stationary, irreducible, reversible Markov chains, Geyer observed that a function Γ*_k_* ≡ γ_2_*_k_* + γ_2_*_k_*_+1_ of the unnormalized fluctuation autocorrelation function γ*_t_* ≡ 〈*a_i_a_i_*+*t*〉 − 〈*a_i_*〉^2^ has a number of pleasant properties (Theorem 3.1 of [15]): It is strictly positive, strictly decreasing, and strictly convex. Some or all of these properties can be exploited to define a family of estimators called *initial sequence methods* (see Section 3.3 of [15] and Section 1.10.2 of [4]), of which the *initial convex sequence* (ICS) estimator is generally agreed to be optimal, if somewhat more complex to implement.^6^

All computations in this manuscript used the fast multiscale method described in Section 5.2 of [10], which we found performed equivalently well to the Geyer estimators (data not shown). This method is related to a multiscale variant of the *initial positive sequence* (IPS) method of Geyer [16], where contributions are accumulated at increasingly longer lag times and the sum of Eq. 11 is truncated when the terms become negative. We have found this method to be both fast and to provide useful estimates of the statistical inefficiency, but it may not perform well for all problems.

## ACKNOWLEDGMENTS

We are grateful to William C. Swope (IBM Almaden Re, search Center) for his illuminating introduction to the use of autocorrelation analysis for the characterization of statistical error, as well as Michael R. Shirts (University of Virginia), David L. Mobley (University of California, Irvine), Michael K. Gilson (University of California, San Diego), Kyle A. Beauchamp (MSKCC), and Robert C. McGibbon (Stanford University) for valuable discussions on this topic, and Joshua L. Adelman (University of Pittsburgh) for helpful feedback and encouragement. We are grateful to Michael K. Gilson (University of California, San Diego), Wei Yang (Florida State University), Sabine Reißer (SISSA, Italy), and the anonymous referees for critical feedback on the manuscript itself. JDC acknowledges a Louis V. Gerstner Young Investigator Award, NIH core grant P30-CA008748, and the Sloan Kettering Institute for funding during the course of this work.

## Footnotes

1 The term *burn-in* comes from the field of electronics, in which a short “burn-in” period is used to ensure that a device is free of faulty components—which often fail quickly—and is operating normally [4].

2 We note that this equality only holds for simulation schemes that sample from the true equilibrium density *π*(*x*), such as Metropolis-Hastings Monte Carlo or Metropolized dynamical integration schemes such as hybrid Monte Carlo (HMC). Molecular dynamics simulations utilizing finite timestep integration without Metropolization will produce averages that may deviate from the true expectation 〈A〉 [2].

3 Note that the *reduced potential* [11] for the isothermal-isobaric ensemble is generally defined as *u^*^*(*x*) = *β*[*u*(*x*) + *pV*(*x*)] to include the pressure-volume term *βpV*(*x*), but in order to demonstrate the performance of this analysis on an observable distinct from the density—which depends on *V*(*x*)—we omit the *βpV*(*x*) term in the present analysis.

4 All Python scripts necessary to reproduce this work—along with data plotted in the published version—are available at: http://github.com/choderalab/automatic-equilibrationdetection

5 available at http://github.com/choderalab/openmmtools

6 Implementations of these methods are provided with the code distributed with this manuscript.

